# Sex-differences in catecholamine transporter expression in the rodent prefrontal cortex following repetitive mild traumatic brain injury and methylphenidate treatment

**DOI:** 10.1101/2025.02.25.640107

**Authors:** Eleni Papadopoulos, Anna Abrimian, Christopher P. Knapp, Jessica A. Loweth, Barry D. Waterhouse, Rachel L. Navarra

## Abstract

Irregular catecholamine transmitter activity is theorized to underly impaired prefrontal cortex (PFC)-mediated executive functions following repetitive mild traumatic brain injury (rmTBI). The psychostimulant, methylphenidate (MPH), enhances catecholamine neurotransmission by blocking reuptake transporters and is used off-label to treat post-TBI executive dysfunction. Although rmTBI and MPH have been shown to independently alter catecholamine transporter levels, the present report evaluated the interactive effects of rmTBI and a sub-chronic therapeutic dose of MPH on expression levels of vesicular monoamine transporter-2 (VMAT2) and norepinephrine reuptake transporter (NET) within subregions of the PFC in both male and female rats. Treatment with MPH restored rmTBI-induced reductions in transporter expression in females. However, in males subjected to rmTBI, MPH exacerbated reductions in transporter expression within the PFC. These results suggest MPH treatment produces beneficial effects in females but exaggerates pathological outcomes in males when used to treat post-rmTBI symptoms.

## Introduction

Traumatic brain injury often impairs prefrontal cortex (PFC)-mediated executive functions, such as attention, working memory, decision making, cognitive flexibility, and inhibitory control over behavior^1^. The PFC contains multiple interconnected regions; the medial prefrontal cortex (mPFC), anterior cingulate cortex (ACC), and orbitofrontal cortex (OFC), that work together to facilitate these processes. Catecholamines, dopamine (DA) and norepinephrine (NE), modulate neural activity within these subregions and thus, influence their operations^2^. Catecholamine concentrations are dysregulated following TBI^3, 4^, suggesting that these imbalances within the PFC may underlie TBI-induced executive dysfunction.

Regulatory proteins, such as those responsible for synthesis, packaging, and clearance of these transmitters, control available catecholamine concentrations. Alterations in these proteins are reported following moderate-severe TBI^5, 6^. However, mild TBI (mTBI) is the most common form of head injury and high-risk populations, such as athletes, military personnel, the elderly, and domestic violence victims are more susceptible to experiencing repeated mTBIs (rmTBI)^7, 8^. Until recently, catecholamine regulatory proteins had not been evaluated following mild or repetitive TBI. With this gap in mind, we demonstrated that alterations in PFC catecholamine regulatory proteins are a plausible mechanism to explain sex differences in executive dysfunction following rmTBI^9, 10^.

Psychostimulant drugs, such as methylphenidate (MPH), block catecholamine reuptake transporters and elevate transmitter concentrations throughout the brain^11^. MPH is often prescribed off-label for TBI due to its efficacy in treating similar symptoms in patients with attention deficit hyperactivity disorder (ADHD), which also arise from catecholamine dysregulation within the PFC^12^. At low therapeutic doses, MPH modifies catecholamine activity with regional specificity for the PFC, which limits side effects driven by subcortical elevations in catecholamine concentrations such as hyperactivity^13^. MPH’s efficacy in treating post-TBI symptoms in clinical and preclinical studies is inconsistent due to limited comprehensive investigations regarding dosage, timing and duration of therapy, and timing and severity of the injuries, especially those categorized as rmTBI’s. Additionally, sex differences in response to drug treatment and injury outcomes are not well established^7, 14^.

Interestingly, chronic administration of low-dose MPH alone influences the regulatory transporters, vesicular monoamine transporter-2 (VMAT2) and norepinephrine transporter (NET), in normal functioning male rodents^15^. However, no reports have assessed potential interactive effects of MPH and injury, or sex, on PFC transporter expression levels. Therefore, we evaluated how rmTBI and low-dose sub-chronic MPH treatment affect transporter expression levels within subregions of the PFC in both male and female rodents.

## Methods

### Animals

Long-Evans rats (Envigo, 17 male and 15 female) were single-housed in a 12h:12h reverse light cycle facility and maintained on a food regulated diet (5g/100g body weight/day) with *ad libitum* water^9, 10^. Animals received sham or impact surgeries in young adulthood, i.e. 9-10 weeks/old. Groups for each sex (male, female) included all combinations of injury (sham, rmTBI) and treatment (saline, 2mg/kg MPH), totaling 8 groups. All procedures adhered to ethical considerations in accordance with the Rowan-Virtua School of Osteopathic Medicine Institutional Animal Care and Use Committee and the National Institutes of Health Guide for the Care and Use of Laboratory Animals.

### Surgeries

The closed head-controlled cortical impact (CH-CCI; Custom Design & Fabrication Inc.) model mimics, in rodents, the functional and biochemical changes of clinical mild TBI cases^9, 10, 16^. CH-CCI was used to induce rmTBI (3 mTBIs within one week, each separated by 2 days) as previously described^10^. Briefly, animals were anesthetized with isoflurane and the skull was exposed by a 2cm midline incision. A 5mm rounded metal impactor tip was aligned with bregma and zeroed along the sagittal suture, then electronically driven at 5.5m/s to a 3.5mm depth below surface with 100ms dwell time.

### Dosing

Methylphenidate hydrochloride (Sigma Aldrich) was dissolved in sterile saline and injected intraperitoneally (i.p.) in 1mL/kg volume. Low-dose MPH (2mg/kg) falls within the therapeutic plasma range in humans (8-40ng/ml) and enhances cognition or reduces impulsivity in various rodent preclinical assays^13, 17-21^. The first dose was administered immediately following the first surgical preparation and continued daily at the same time for 9 days, concluding 48hr post-final preparation.

### Western Blotting

One hour post-final dose, animals were anesthetized and the mPFC, ACC, and OFC^22^ were dissected to determine protein expression levels of VMAT2 (responsible for packaging catecholamines into vesicles for subsequent release at the synapse) and NET (responsible for catecholamine uptake and synaptic clearance). The dopamine transporter (DAT) takes up DA and is also blocked by MPH, but it is sparsely expressed within the PFC and therefore, wasn’t evaluated in this study^23^. As previously described^9, 10^, protein (15 ug/lane) from the collected tissue was electrophoresed, transferred to polyvinylidene difluoride membranes (Bio-Rad), and probed with rabbit anti-VMAT2 (1:1000; Abcam) or anti-NET (1:1000; Abcam) primary antibody followed by goat anti-rabbit secondary antibody conjugated with peroxidase (1:10,000; Rockland Immunochemicals, Inc.). β-actin (1:2000; MilliporeSigma) was the loading control. Blots were imaged using Azure c400 Biosystems imaging system and analyzed using AzureSpot Analysis Software (Azure Biosystems).

### Statistical analysis

Analysis was performed using GraphPad Prism software. Males and females were analyzed separately using an ordinary two-way ANOVA with injury and treatment conditions as between-subject factors. Dunnett’s multiple comparisons tests were used to compare individual group differences to the sham/saline control group when a main effect or interaction was found. Statistical significance was determined by a p value < 0.05.

## Results

### RmTBI and MPH effects on VMAT2 expression levels

*VMAT2 expression levels in males*. Analysis of VMAT2 protein expression levels within the mPFC revealed a main effect of injury [F (1, 13) = 11.70, p = 0.0046], but no significant effects of treatment [F (1, 13) = 1.807, p = 0.2018] nor an injury-by-treatment interaction [F (1, 13) = 1.029, p = 0.3289] (Fig. 1A). Dunnett’s multiple comparisons revealed a reduction in VMAT2 levels within the rmTBI/saline group (p = 0.0236) and further decrease in the rmTBI/MPH group (p = 0.0110) suggesting that MPH treatment may exacerbate rmTBI-induced decreases in VMAT2 expression. In the ACC, there was a main effect of injury [F (1, 13) = 9.712, p = 0.0082], but not treatment [F (1, 13) = 3.646, p = 0.0785] or an interaction [F (1, 13) = 0.1921, p = 0.6683] (Fig. 1B). Multiple comparisons revealed a near-significant decrease in VMAT2 levels in the rmTBI/saline group (p = 0.0717), while the rmTBI/MPH group demonstrated a highly significant decrease (p = 0.0077). In the OFC, there was a treatment effect [F (1, 13) = 6.723, p = 0.0223], but no effect of injury [F (1, 13) = 1.317, p = 0.2718] or interaction [F (1, 13) = 0.3616, p = 0.5579] (Fig. 1C). Multiple comparisons revealed a decrease in VMAT2 within the rmTBI/MPH group only (p = 0.0441). Although rmTBI alone was not sufficient to significantly decrease VMAT2 expression in the ACC and OFC, the combination of rmTBI/MPH produced noticeable decreases in all 3 subregions suggesting these two factors played complimentary roles to decrease VMAT2 in males.

**Figure 1.**
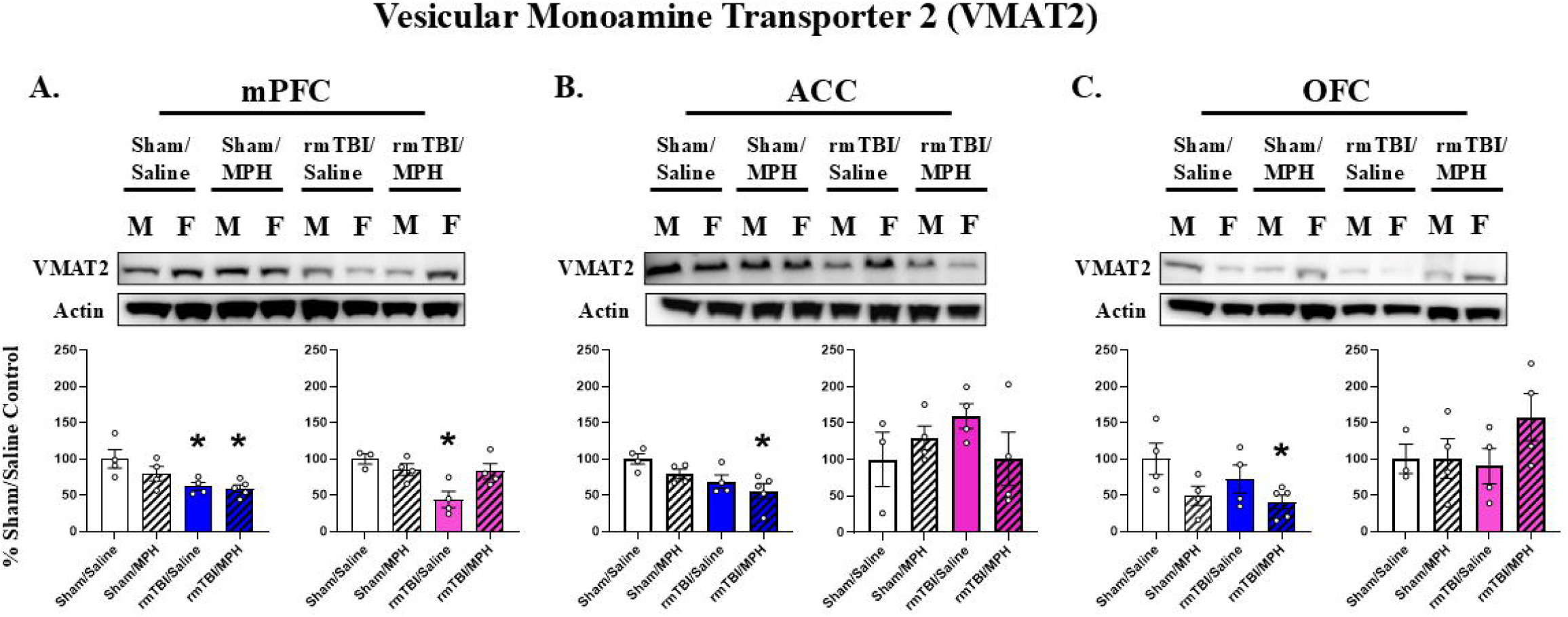
Protein expression levels of VMAT2 within the A) mPFC, B) ACC, and C) OFC following rmTBI and MPH treatment. Graphs represent mean percent change in total protein levels ±SEM as compared to sham/saline controls 48-hr post-final surgery and 1-hr post-final administration of saline or MPH (2mg/kg, i.p., hashed bars) for males (left, blue) and females (right, pink). Note that rmTBI-induced reductions of VMAT2 expression levels within the mPFC, ACC and OFC subregions are either unaffected or exacerbated by MPH administration in males but reversed within the mPFC of females. *denotes p < 0.05 from Sham/Saline analyzed with Two-Way ANOVA and Dunnett’s Multiple Comparisons Tests. N = 3-5 per group. Abbreviations: VMAT2 (vesicular monoamine transporter-2), mPFC (medial prefrontal cortex), ACC (anterior cingulate cortex), OFC (orbitofrontal cortex), MPH (methylphenidate).

*VMAT2 expression levels in females*. Analysis of mPFC VMAT2 expression levels revealed a main injury effect [F (1, 11) = 8.982, p = 0.0121] and injury-by-treatment interaction [F (1, 11) = 7.425, p = 0.0198], but no effect of treatment [F (1, 11) = 1.526, p = 0.2425] (Fig. 1A). Importantly, multiple comparisons revealed a decrease in VMAT2 levels within the rmTBI/saline group (p = 0.0063), that was not observed following rmTBI/MPH (p = 0.4958), suggesting this MPH treatment regimen prevented the rmTBI-induced decrease in female mPFC VMAT2. No effects of injury, treatment, or interaction were observed on VMAT2 levels in ACC [F (1, 11) = 0.3132, p = 0.5869; F (1, 11) = 0.2782, p = 0.6084; F (1, 11) = 2.501, p = 0.1420, respectively] (Fig. 1B) or OFC [F (1, 11) = 0.7238, p = 0.4131; F (1, 11) = 1.507, p = 0.2452; F (1, 11) = 1.475, p = 0.2500, respectively] (Fig. 1C).

### RmTBI and MPH effects on NET expression levels

*NET expression levels in males*. Analysis of mPFC NET protein expression levels revealed no significant effects of treatment [F (1, 13) = 0.8616, p = 0.3702] nor an interaction [F (1, 13) = 1.901, p = 0.1913, respectively] (Fig. 2A). However, with a trend towards an injury effect [F (1, 13) = 3.304, p = 0.0922], the data clearly demonstrates a 32.3% reduction following rmTBI/saline and 27.7% reduction following rmTBI/MPH compared to sham/saline. Although shy of significance, the patterns of rmTBI-induced transporter reductions with no benefit from MPH administration are comparable to those observed in male mPFC VMAT2. No treatment, injury, or interaction effects were detected in ACC NET [F (1, 12) = 0.01855, p = 0.8939; F (1, 12) = 0.8607, p = 0.3718; F (1, 12) = 0.06122, p = 0.8088, respectively] (Fig. 2B). In the OFC (Fig. 2C), there was no effect of injury [F (1, 13) = 1.375, p = 0.2621] or interaction [F (1, 13) = 0.2312, p = 0.6386], but there was a treatment effect [F (1, 13) = 7.268, p = 0.0183] with a decrease in OFC NET levels following rmTBI/MPH (p = 0.0372) only (Fig. 2C). Once more suggesting the rmTBI/MPH interaction is necessary to produce significant decreases as observed with male OFC VMAT2 (Fig. 1C).

**Figure 2.**
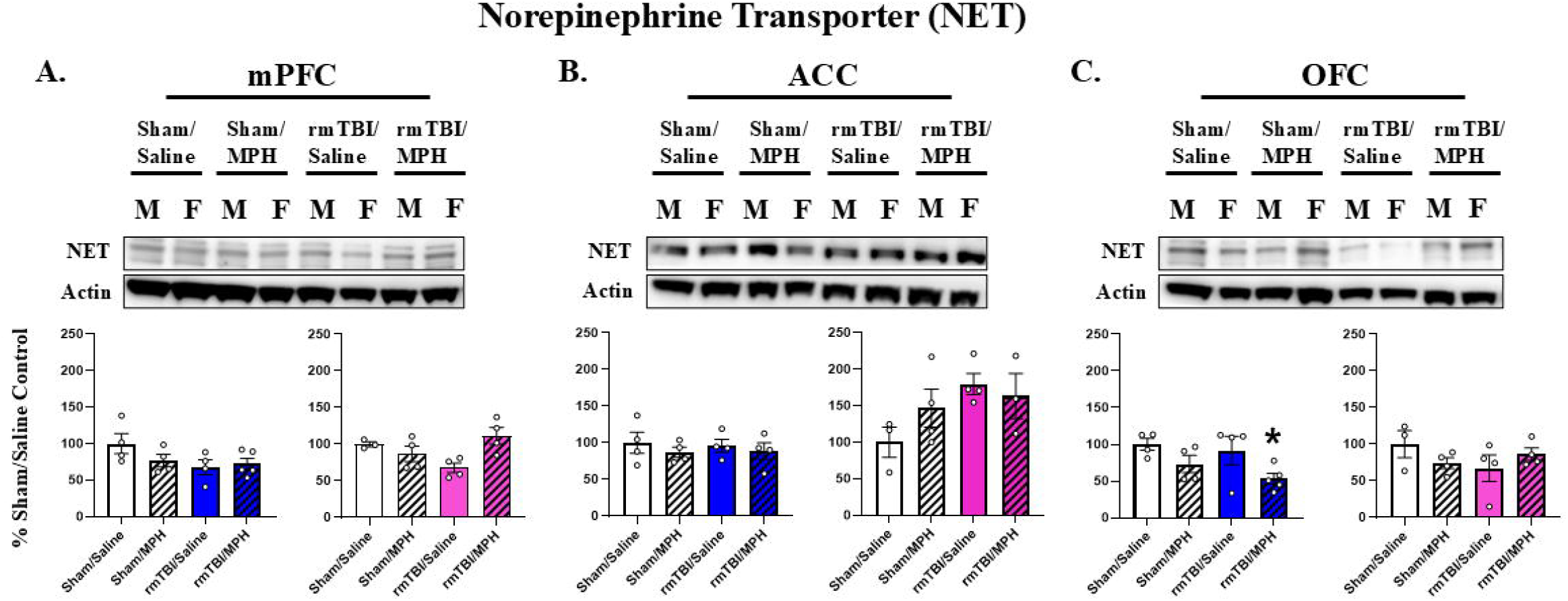
Protein expression levels of NET within the A) mPFC, B) ACC, and C) OFC following rmTBI and MPH treatment. Graphs represent mean percent change in total protein levels ±SEM as compared to sham/saline controls 48-hr post-final surgery and 1-hr post-final administration of saline or MPH (2mg/kg, i.p., hashed bars) for males (left, blue) and females (right, pink). Note that mPFC NET displays similar patterns of protein expression as compared to mPFC VMAT2 (Fig 1). Only the combination of rmTBI and MPH administration significantly decreased OFC NET in males. *denotes p < 0.05 from Sham/Saline analyzed with Two-Way ANOVA and Dunnett’s Multiple Comparisons Tests. N = 3-5 per group. Abbreviations: NET (norepinephrine transporter), mPFC (medial prefrontal cortex), ACC (anterior cingulate cortex), OFC (orbitofrontal cortex), MPH (methylphenidate).

*NET expression levels in females*. There was a significant injury-by-treatment interaction [F (1, 11) = 9.388, p = 0.0108] on female mPFC NET, but no effects of injury [F (1, 11) = 0.1593, p = 0.6975] or treatment [F (1, 11) = 2.642, p = 0.1324] alone (Fig 2A). Although no significant changes were detected by multiple comparisons, there was a strong 32% decrease in NET in the rmTBI/saline group (p = 0.0881) compared to sham/saline. This seems to be prevented by treatment in the rmTBI/MPH group (p = 0.7286), following the same pattern as female mPFC VMAT2 (Fig. 1A). No effects of injury, treatment, or interaction were observed on NET levels in ACC [F (1, 10) = 4.209, p = 0.0673; F (1, 10) = 0.4105, p = 0.5361; F (1, 10) = 1.813, p = 0.2078, respectively] (Fig. 2B) or OFC [F (1, 11) = 0.5642, p = 0.4683; F (1, 11) = 0.06179, p = 0.8083; F (1, 11) = 2.748, p = 0.1256, respectively] (Fig. 2C).

## Discussion

The present results demonstrate rmTBI and a sub-chronic low dose of MPH display interactive effects on catecholamine transporter expression within subregions of the PFC in a sex-dependent manner. Our findings suggest MPH treatment produces beneficial effects in females but exaggerates protein-level perturbations in males on following rmTBI. Additionally, we are the first to report VMAT2 expression changes in the PFC following a closed-head rodent model of rmTBI.

In the mPFC, rmTBI reduced expression levels of VMAT2 in both sexes. MPH treatment reestablished VMAT2 levels comparable to sham/saline following rmTBI in females, but not males. Although not statistically significant given the conservative requirements of multiple comparisons, this pattern of non-specific rmTBI-induced reduction but female-specific MPH-induced restoration was also observed with mPFC NET following both manipulations. Together, decreased transporter levels following rmTBI suggest a compensatory response to low catecholamine levels following TBI, as previously reported^3^. In an injury-induced hypo-catecholaminergic state, decreased transmitter concentrations signal less substrate availability and therefore, less need for reuptake and repackaging following injury, which leads to downregulation of the transporters required to mediate these actions^24^. However, when NET is blocked with MPH, higher concentrations of the remaining catecholamines are left within the synapse and available for transport. Eventually, rising levels of extracellular transmitter would again require reuptake and repackaging, signaling need to upregulate NET and VMAT2, respectively, as demonstrated in the mPFC of females following the combination of both rmTBI and MPH.

Interestingly, in males only the combination of rmTBI and MPH was sufficient to significantly decrease VMAT2 expression levels in the ACC and OFC, as well as NET expression in the OFC. Although we reported decreases in PFC NET expression following a similar rmTBI protocol^9^, this potentiated decrease was surprising in lieu of findings demonstrating that similar MPH regimens increased VMAT2 and NET expression levels within the frontal cortex of uninjured male rodents^15^, leading expectations that MPH would reverse rmTBI-induced transporter reductions as demonstrated in our females. Nevertheless, we reveal the potential for MPH to exacerbate rmTBI-induced biochemical signatures that may underlie further impairments of executive function in males, especially during this critical time point of young adult PFC development.

MPH’s opposing sex-specific effects on transporter expression following injury could be due to differential pharmacokinetics and circulating drug concentrations in males vs females^25^. For example, sex-specific MPH-induced influences in performance and drug sensitivity have been reported, where a higher MPH dosing regimen improved injury-induced spatial memory deficits in males but increased locomotor activity in females, respectively^14^. Therefore, it was unsurprising that the lower dose of MPH used in this study was sufficient for females, but males may require a higher dose to achieve the same restorative effects. Indeed, the observed sex differences could reflect the mere fact that the female brain is under constant hormonal flux, providing an additional layer of influence over responses to rmTBI and drug-induced alterations of transmitter concentrations and protein expression levels.

Overall, these findings demonstrate that regulation of PFC transporter expression is dynamic and adaptive in response to experimental injury and drug manipulations that perturb catecholamine homeostasis. Although these interactive effects on transporter expression are a glimpse into the catecholaminergic regulatory processes, the present report suggests that this dosing regimen of MPH following rmTBI produces beneficial effects in females, but detrimental effects in males. Future directions will investigate other key proteins involved in catecholamine regulation and behavioral outcomes using this dosing regimen of MPH following rmTBI. To conclude, we believe this work is both timely and highlights the importance of considering sex-dependent treatment strategies associated with off-label use of MPH for treating post-TBI executive dysfunction.

## Acknowledgements

The authors thank Douglas P. Fox and Karen L. Joyce for their technical assistance in this study.

## Data availability statement

The data underlying this article will be shared on reasonable request to the corresponding author.

